# Development and validation of a genome-informed multiplex PCR for specific detection of typhoidal *Salmonella* serovars

**DOI:** 10.64898/2026.06.01.729223

**Authors:** Jobin John Jacob, Praveen Thilagan, Pavithra Sathya Narayanan, KB Santhosh, R Subbulakshmi, Aravind Velmurugan, Monisha Priya Teekaraman, Nirmaladevi Ponnusamy, Ayyan Raj Neeravi, Jacob John, Kamini Walia, Balaji Veeraraghavan

## Abstract

Enteric fever caused by *Salmonella enterica* serovars Typhi and Paratyphi A, B and C remains a major public health burden in endemic regions. Existing molecular assays frequently demonstrate limited specificity due to cross-reactivity with non-typhoidal *Salmonella* (NTS). In this study, we developed and validated a genomics-informed multiplex PCR assay capable of simultaneously differentiating all four typhoidal *Salmonella* serovars. A curated dataset of 3,239 Salmonella genomes, including *S*. Typhi (n=361), *S*. Paratyphi A (n=453), *S*. Paratyphi B (n=511), *S*. Paratyphi C (n=62), and NTS genomes (n=1,853), was used for comparative genomic analysis. Thirty published PCR targets were evaluated *in silico*, followed by pangenome and SNP analyses to identify discriminatory loci for mismatch amplification mutation assay (MAMA)-based primer design. Candidate primers were validated using *in silico* PCR, BLASTn analysis, and laboratory testing against a panel of typhoidal Salmonella, clinical NTS isolates, and non-Salmonella bacterial pathogens. *In silico* evaluation demonstrated substantial cross-reactivity among many published targets, whereas SNP-informed primer design targeting *staG* (*S*. Typhi), SPA0152 (*S*. Paratyphi A), SPAB_03490 (*S*. Paratyphi B), and SPC_0571 (*S*. Paratyphi C) achieved predicted specificities of 98–100% while retaining high analytical sensitivity (>97%) across target genomes. Combined with a pan-Salmonella *invA* target, the multiplex assay precisely identified all target serovars *in vitro* with minimal cross-reactivity. These findings demonstrate that genomics-informed SNP-based primer design enables reliable multiplex differentiation of typhoidal *Salmonella* serovars and provides a scalable framework for improving enteric fever diagnosis and surveillance in endemic settings.

**Importance:** Typhoidal *Salmonella* serovars remain major causes of enteric fever in endemic regions, yet molecular differentiation from non-typhoidal *Salmonella* (NTS) remains challenging because of extensive genomic conservation and cross-reactivity of commonly used diagnostic targets. In this study, we combined large-scale comparative genomics of 3,239 *Salmonella* genomes with SNP-informed primer design to develop a multiplex PCR assay capable of simultaneously differentiating all four typhoidal serovars (*S*. Typhi, *S*. Paratyphi A, B, and C) from NTS and other non-Salmonella pathogens. Unlike conventional gene-content-based assays, this approach incorporated lineage-specific SNPs and mismatch amplification strategies to improve specificity while maintaining high analytical sensitivity. *In silico* evaluation demonstrated high diagnostic performance across diverse global lineages, while *in vitro* testing confirmed accurate serovar-level discrimination with minimal cross-reactivity. These findings demonstrate the value of population-scale genomics for molecular assay development and provide a scalable framework for improving diagnosis and surveillance of enteric fever in endemic settings.

## Introduction

Enteric fever, primarily caused by *Salmonella enterica* serovars Typhi, *S*. Paratyphi A and, to a lesser extent, by serovars *S*. Paratyphi B, and C, continues to pose a substantial public health challenge, particularly in low- and middle-income countries (1). Despite the introduction of vaccines and improvements in food safety, water quality, hygiene, and sanitation, typhoid fever remains a major threat to global health. The burden is especially high in resource-limited regions of South Asia, Southeast Asia, sub-Saharan Africa, and Oceania (2). A recent global estimate reported a 62.1% decline in typhoid fever and a comparable decline in paratyphoid fever from 1990 to 2021 (3). Nevertheless, rapid urbanization, population mobility, and increased international travel continue to create conditions that sustain transmission and increase the risk of wider global spread (4).

Early and accurate diagnosis of enteric fever is critical, as delays or inappropriate therapy can lead to severe acute complications, chronic carriage, and increased transmission (5). Clinical diagnosis alone is unreliable, particularly in endemic settings as presenting symptoms including prolonged fever, headache, malaise, and abdominal discomfort overlap with those of other common infections such as malaria, dengue, and brucellosis (6). Blood culture remains the reference standard for diagnosis, but it is constrained by a turnaround time (2–4 days) a modest sensitivity of ranging from 40% to 87% and limited availability in many resource-limited settings (7,8). Widely used serological tests such as the Widal test suffer from poor sensitivity and specificity in endemic regions, while newer rapid serological assays (e.g., Tubex TF, Typhidot IgG/IgM) offer only incremental improvements and cannot replace culture-based diagnosis (9).

Molecular diagnostic approaches including conventional PCR, quantitative PCR (qPCR), and loop-mediated isothermal amplification (LAMP) have demonstrated promise for the detection of typhoidal *Salmonella* (10,11,12,13). However, many existing assays exhibit variable performance and face significant challenges in endemic settings, particularly due to cross-reactivity with closely related non-typhoidal *Salmonella* (NTS) serovars (12). Most of the commonly used marker genes for typhoidal *Salmonella* detection, including *tviB* and *staG*, were characterized and adopted for diagnostics prior to the widespread availability of NGS data. To enhance specificity, multiple genetic targets have been evaluated for the detection of *S*. Typhi and *S*. Paratyphi, including *tviB, staG, stoD*, STY1076, STY0307, and STY1599 for *S*. Typhi, and *fliC-*a and SPA2308 for *S*. Paratyphi A (10). Nevertheless, several of these targets share homologous sequences with NTS or other members of the Enterobacterales, limiting diagnostic accuracy (14). In this study, we report the development and validation of a conventional multiplex PCR assay capable of simultaneously detecting four typhoidal *Salmonella* serovars offering a more accurate diagnostic tool to improve clinical management and treatment of enteric fever in endemic settings.

## Methods

### Retrieval and curation of *Salmonella* genome sequencing data

Whole-genome sequencing (WGS) data for contemporary clinical isolates of *Salmonella* Typhi, Paratyphi A, B, and C predominantly generated at the study centre as part of prior genomic surveillance were retrieved from the European Nucleotide Archive (ENA; https://www.ebi.ac.uk/ena). In addition, a representative global collection of genomes encompassing both typhoidal and NTS serovars was obtained from EnteroBase (https://enterobase.warwick.ac.uk/species/index/senterica). Genomes were selected based on sequence types (STs) to capture broad temporal diversity within each serovar. Genome assemblies that did not meet quality criteria were excluded from downstream analyses. The final curated dataset, comprising 3239 genomes, was used as the reference panel for *in silico* evaluation of PCR primers and for the identification of candidate diagnostic targets (**Table S1**).

### *In silico* evaluation of existing PCR targets and primers

Previously published PCR primers and gene targets commonly used for the detection of *Salmonella* Typhi, Paratyphi A, B, and C were compiled from the literature (**Table S2**). *In silico* PCR analyses were performed using the Perl script in_silico_pcr.pl (https://github.com/egonozer/in_silico_pcr) against a curated genome dataset. The analysis was used to evaluate primer specificity, predicted amplification profiles, amplicon generation efficiency, and amplicon size distribution across typhoidal and non-typhoidal Salmonella serovars. Primer–target matching was assessed using a stringency threshold of ≤2 total mismatches. Accordingly, *in silico* amplification events with ≤2 mismatches were considered as potential positive signals. Sensitivity and specificity were calculated from the curated dataset (n = 3,239 genomes) based on counts of true positives (TP), true negatives (TN), false positives (FP), and false negatives (FN).

To further validate target specificity, reference gene sequences corresponding to each PCR target were analysed using National Center for Biotechnology Information Basic Local Alignment Search Tool nucleotide (NCBI BLASTn) against publicly available databases to confirm target identity and assess their presence in non-target serovars. Primer–genome alignments were further examined to identify potential cross-reactivity and false-positive amplification events.

### Pangenome & SNP analysis and identification of serovar-specific targets

To identify novel serovar-specific diagnostic targets, pangenome analysis was performed on the curated genome dataset. Annotated genome assemblies (GFF3 files), generated using Bakta v1.12, (https://github.com/oschwengers/bakta) were used as input for pangenome analysis with Panaroo v1.5.2 (https://github.com/gtonkinhill/panaroo) with default parameters. The resulting gene presence–absence matrix was grouped into five predefined categories *S*. Typhi, Paratyphi A, B, C, and NTS and clustered using the Twilight analysis package (https://github.com/ghoresh11/twilight) with default thresholds. Comparative analysis of core and accessory gene content was performed to identify genes uniquely conserved within individual typhoidal serovars and absent from NTS.

In parallel, single nucleotide polymorphism (SNP) analysis was conducted using the Snippy v4.6.0 variant calling pipeline (https://github.com/tseemann/snippy). Genomes from the curated dataset were mapped against the annotated reference genome *Salmonella* Typhimurium LT2 (NC_003197.2). Custom-written Bash scripts were used to extract serovar-specific patterns of mutation accumulation across the dataset. Genes harbouring conserved serovar-specific missense or frameshift SNPs were shortlisted as candidates for mismatch-based discriminatory PCR primer design.

### Primer design based on unique genes and serovar-specific SNPs

Primers were designed based on findings from pangenome and SNP analyses. Where serovar-specific genes were identified, these loci were prioritised for primer design. For *S*. Typhi and *S*. Paratyphi C candidate targets demonstrating complete presence across all genomes were selected to ensure maximal analytical sensitivity. As some of these loci were also present in non-typhoidal *Salmonella*, serovar specificity was enhanced by incorporating target conserved sequence variants identified through SNP analysis into primer binding regions, enabling mismatch-based discrimination of non-target serovars.

In instances where unique genes were not available, conserved loci containing discriminatory SNPs were selected for mismatch amplification mutation assay (MAMA) - PCR design using NCBI Primer-BLAST (https://www.ncbi.nlm.nih.gov/tools/primer-blast/). For SNP-based targets, discriminatory nucleotides were positioned at the 3′ end of primers to enhance allelic specificity. All primer sets were evaluated *in silico* for melting temperature compatibility, GC content, and the absence of secondary structures formation. Potential cross-reactivity was assessed by *in silico* PCR analysis (http://insilico.ehu.eus/PCR/index.php?mo=Salmonella), followed by BLASTn analysis of predicted amplicon sequences against the NCBI nucleotide database, performed both including and excluding the target serovar to confirm specificity.

### Experimental validation of multiplex PCR primers

Designed primers were synthesized and initially evaluated in singleplex PCR assays using a panel of well-characterized reference strains and clinical isolates representing typhoidal *Salmonella* (n = 5), non-typhoidal *Salmonella* (NTS; n = 17), and non-*Salmonella* bacterial pathogens (n = 8). Genomic DNA was extracted from single bacterial colonies grown on culture plates using the QIAamp DNA Mini Kit (Qiagen Hilden, Germany) according to the manufacturer’s instructions, followed by DNA quantification and quality assessment prior to PCR analysis. Primer sets demonstrating optimal amplification efficiency and target specificity were subsequently combined into a multiplex PCR assay. Multiplex PCR reactions were performed using the Qiagen Multiplex PCR kit (Qiagen, Hilden, Germany) on an ProFlex thermal cyclers (Applied Biosystems, Waltham, MA, USA) in a final reaction volume of 20 µL containing 10 µL of 2× multiplex master mix, 2 µL of Q-solution, 2 µL of primer mix, 4 µL of nuclease-free water, and 2 µL of template DNA. Primers targeting *invA* and *staG* were used at a final working concentration of 10 µM, whereas primers targeting SPA0152, SPAB_03490, and SPC0571 were used at 5 µM. Thermal cycling conditions consisted of an initial denaturation at 95°C for 10 min, followed by 30 cycles of denaturation at 94°C for 30 s, annealing at 58°C for 1 min 30 s, and extension at 72°C for 1 min 30 s, with a final extension at 72°C for 10 min. Amplified products were resolved by electrophoresis on 2% agarose gels at 120 V for 50 min.

### Analytical performance of the multiplex PCR assay

The analytical performance of the multiplex PCR assay was evaluated using the same panel of reference strains and clinical isolates. Sensitivity and specificity were calculated based on the number of true positives (TP), true negatives (TN), false positives (FP), and false negatives (FN) obtained from the multiplex PCR results in comparison with reference identification methods. Sensitivity was calculated as: Sensitivity = TP / (TP + FN) and specificity was calculated as: Specificity = TN / (TN + FP).

## Results

### Composition and genetic diversity of the curated *Salmonella* genome dataset

The curated genome dataset comprised 3,239 *Salmonella* genomes, including 361 (11.1%) *S*. Typhi, 453 (13.9%) *S*. Paratyphi A, 511 (15.7%) *S*. Paratyphi B, 62 (2%) *S*. Paratyphi C, and 1,853 (57.2%) non-typhoidal *Salmonella* (NTS) genomes (**Table S1**). The collection included both contemporary clinical isolates generated at the study centre and representative global genomes retrieved from EnteroBase. MLST analysis demonstrated substantial diversity within each serovar. *S*. Typhi isolates were dominated by globally disseminated ST1/ST2-associated lineages, whereas *S*. Paratyphi A isolates were largely represented by ST85 and ST129. *S*. Paratyphi B and C displayed greater lineage diversity, while the NTS collection comprised 1,342 sequence types across 626 serovars. cgMLST phylogeny generated using EnteroBase and visualized with GrapeTree demonstrated broad phylogenetic diversity and clear separation between typhoidal and NTS serovars (**Figure 1**), supporting the suitability of the dataset for comprehensive *in silico* evaluation of diagnostic targets.

**Figure 1:**
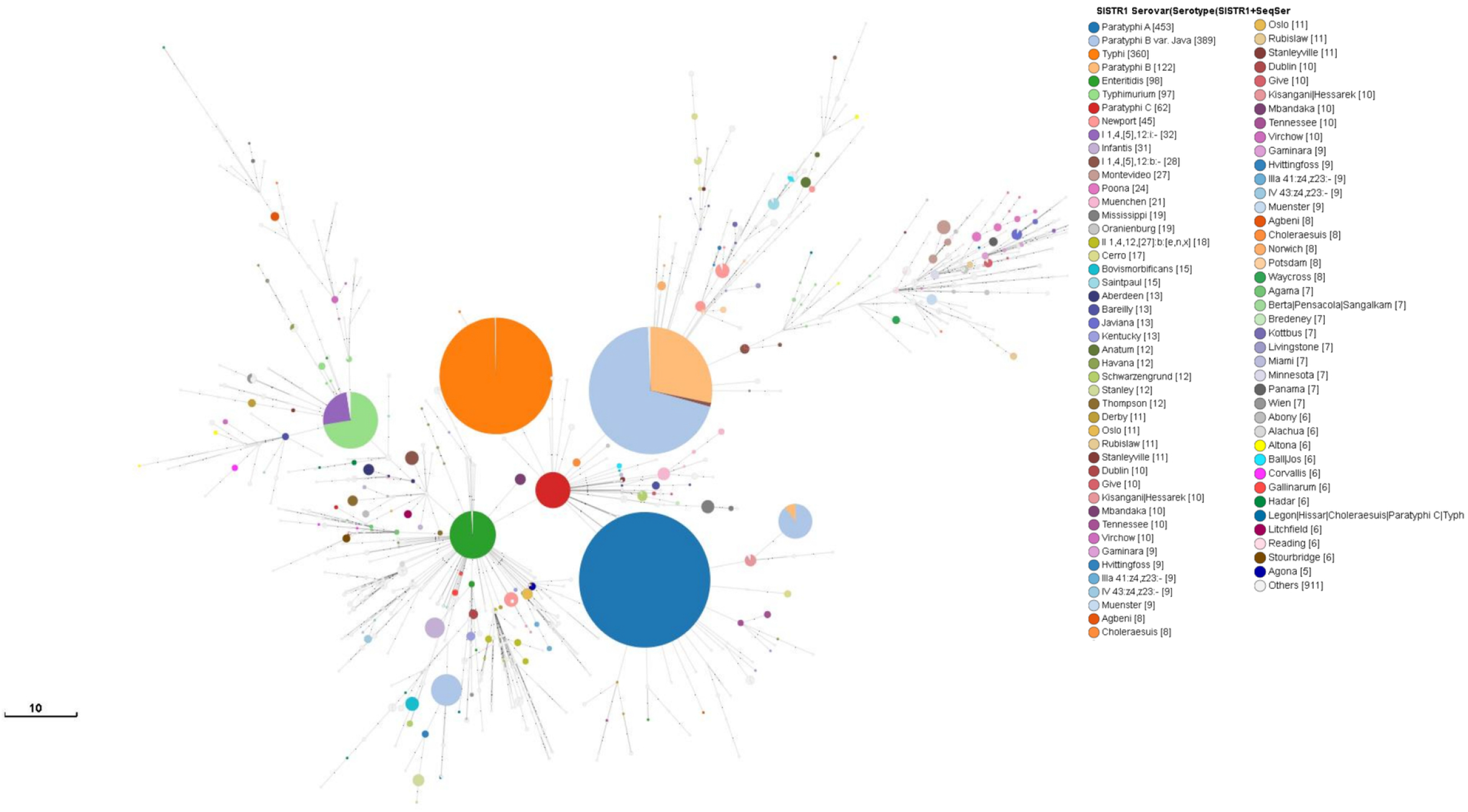
Minimum spanning tree (MST) showing the population structure of the curated Salmonella genome dataset (n = 3,239) based on core genome multilocus sequence typing (cgMLST) available at Enterobase. The MST was generated from cgMLST allelic profiles and visualized using GrapeTree v1.5.0. Each node represents a unique sequence type (ST) or closely related clonal group, with node size proportional to the number of genomes represented. Nodes are coloured according to serovar assignment. Tree nodes were positioned through dynamic rendering, and node style was adjusted by fine-tuning node size and kurtosis for improved visualization. Branch lengths correspond to the number of allelic differences between connected nodes, with the scale bar representing 10 allelic differences.

### *In silico* assessment of existing PCR primers for Typhoidal Salmonella

*In silico* evaluation of 30 previously published PCR primers demonstrated substantial variability in diagnostic performance across typhoidal *Salmonella* serovars (**Table S2**). For *S*. Typhi, several commonly used targets, including *staG* (STY0201), *tviB* (STY4661), *fliC-*d (STY2167), STY0307 and multiple other loci, demonstrated consistently high predicted sensitivity (≥98%) across lineages but variable specificity due to cross-reactivity with NTS serovars. Among evaluated targets, STY1599 demonstrated the best overall performance with 99.45% sensitivity and 99.97% specificity. A similar pattern was observed for *S*. Paratyphi A, where targets such as SPA2308, SPA0180, SPA2473, SPA2539, and *fliC-a* (SPA0911) accurately identified most *S*. Paratyphi A genomes (sensitivity ≥99%) but demonstrated considerable cross-reactivity with non-target serovars, with specificity dropping as low as 81.9% for certain loci (**Table S2**).

For *S*. Paratyphi B, diagnostic performance varied considerably by lineage and primer strategy. Combination targets such as *sseJ*–/*srfJ*+ used for detection of *S*. Paratyphi B PG1 (var. *sensu stricto*) and *sseJ*+/*srfJ*+ for *S*. Paratyphi B var. Java, achieved high sensitivity (100%) and specificity (98.08%) for PG1 isolates. Additional combination targets, including *rfbJ* + Hb, showed poor and lineage-dependent performance. (**Table S2**). The marker STM3356, demonstrated perfect sensitivity but poor specificity (9.06%), limiting standalone utility. In contrast, SPAB_01124 showed high sensitivity (90.75%) and specificity (98.17%) but lacked discriminatory ability between PG1 and var. Java lineages. For *S*. Paratyphi C, SPC4629 achieved 97.22% sensitivity and 99.75% specificity, whereas SPC0869 showed lower sensitivity (83.56%) but retained high specificity (99.72%). Collectively, these analyses highlighted major limitations in existing assays and emphasized the need for genomics-informed serovar-specific targets.

### Pangenome analysis

Pan-genome analysis of the curated *Salmonella* genome dataset (n = 3,239) using Panaroo identified a total of 48,574 gene clusters. Of these, 3,002 genes (6.18%) were classified as core genes present in ≥99% of genomes, and 400 genes (0.8%) were designated as soft-core (95– 99%). The shell genome (15–95%) comprised 1,521 genes (3.13%), while the majority of gene clusters (43,651 genes; 89.86%) were classified as cloud genes (0–15%), indicating extensive accessory gene diversity across the dataset. The relative contributions of core, soft-core, shell, and cloud gene fractions are shown in **Figure S1a**. To identify serovar-specific loci, genomes were stratified into *S*. Typhi, *S*. Paratyphi A (SPA), *S*. Paratyphi B (SPB), *S*. Paratyphi C (SPC), and NTS groups. Lineage-specific core gene analysis identified 25 candidate genes in *S*. Typhi, 11 in SPA, and two in SPC, whereas no lineage-specific core genes were detected for SPB or NTS (**Figure S1b; Table S3**).

Subsequent BLASTn screening demonstrated marked differences in target specificity and diagnostic suitability. Among the 25 putative *S*. Typhi-specific genes, 12 were highly conserved (>99%) and minimally represented (<0.1%) in non-target serovars. However, all gene candidates except the Vi antigen represented pseudogenes misannotated as unique loci by Bakta/Panaroo and were excluded. Similarly, all putative SPA-specific genes were identified as misannotated pseudogenes. In contrast, the two SPC-associated genes retained partial discriminatory potential and were shortlisted for further evaluation (**Table S3**).

### Primer design and *in silico* validation

Following comparative genomic analysis, the *staG, stoD*, and STY1599 loci were selected as candidate targets for *S*. Typhi primer development based on their widespread use in diagnostic assays and near-universal presence across the curated *S*. Typhi genome dataset. Although *staG* and *stoD* were not unique at the gene-content level, comparative analysis identified conserved *S*. Typhi-specific SNPs absent in NTS serovars. Primers incorporating discriminatory 3′ mismatches retained predicted amplification across 100% of *S*. Typhi genomes while eliminating amplification in non-target serovars such as *S*. Napoli, *S*. Moero, and *S*. Liverpool (**Figure S2a**). Optimized *stoD* and STY1599 primers achieved sensitivities/specificities of 99.24%/98.82% and 99.45%/100%, respectively (**Table 1**). Despite strong performance, *staG* was selected for multiplex development due to its superior combined sensitivity and specificity.

**Table 1:**
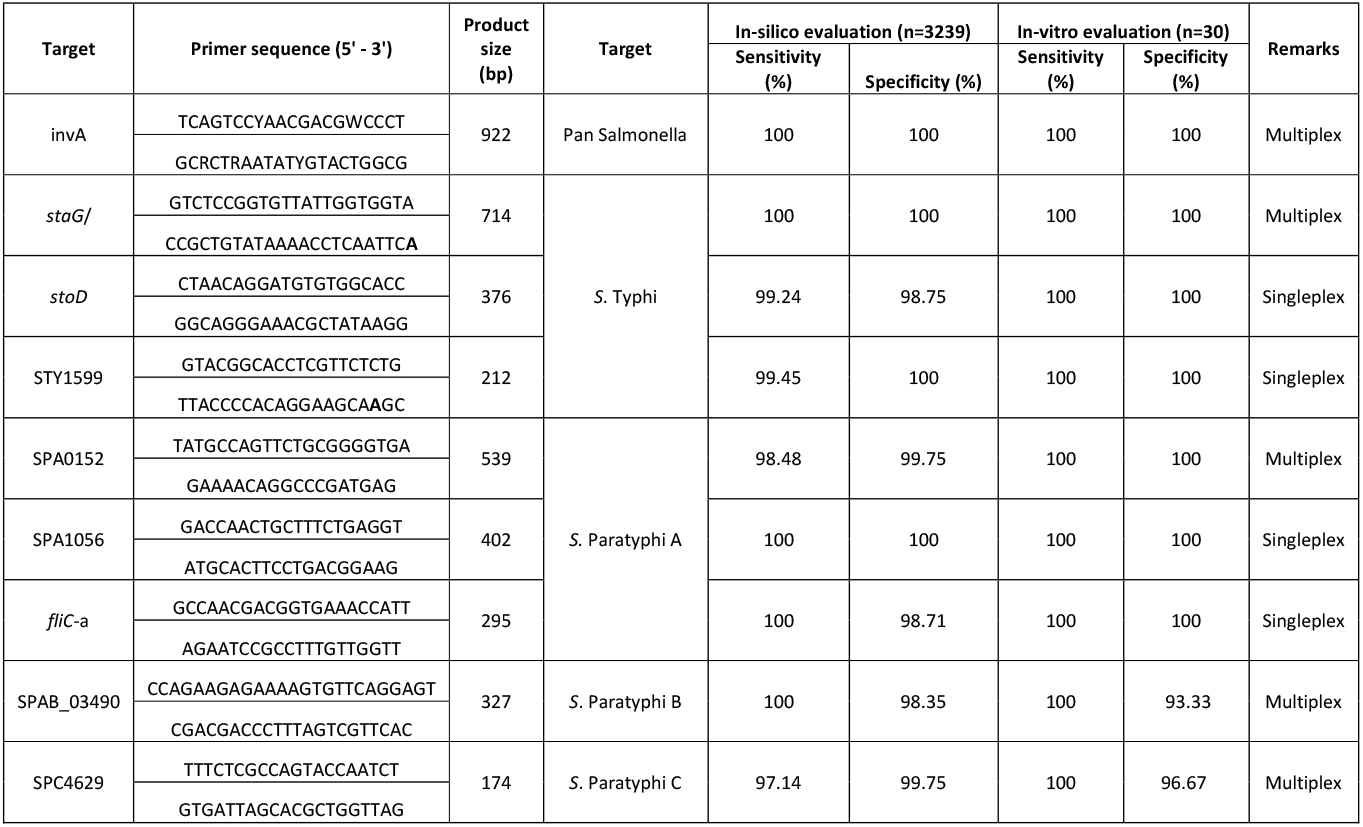
Primer sequences and analytical performance of multiplex and singleplex PCR targets for detection of typhoidal Salmonella serovars.

For *S*. Paratyphi A, pangenome-derived candidate genes were largely identified as pseudogenes, necessitating SNP-based primer design targeting *fliC-a*, SPA0152, and SPA0156. MAMA-based primers incorporating deliberate 3′ mismatches yielded predicted amplicons of 295 bp (*fliC-a*), 539 bp (SPA0152), and 402 bp (SPA0156), respectively (**Table 1; Figure S2b**). *In silico* performance evaluation showed that primers targeting *fliC-a* demonstrated 100% sensitivity and 98.71% specificity, SPA0152 showed 98.48% sensitivity and 99.75% specificity, whereas SPA0156 achieved 100% sensitivity and specificity. Despite the superior analytical performance of SPA0156, the SPA0152 locus was selected for inclusion in the multiplex panel due to its larger amplicon size (539 bp), which provided improved band separation and resolution relative to other targets in the multiplex gel-based assay.

For *S*. Paratyphi B, pangenome and SNP analyses did not identify any single serovar-defining genes or mutation that was completely absent from NTS serovars. A conserved missense SNP within the SPAB_03490 (STM2810) locus was selected for primer design, as it was present across all *S*. Paratyphi B (including both Java and PG1 lineages) genomes but was also detected in approximately 3% of rare NTS serovars. In this case, primer specificity relied directly on the serovar-associated missense SNP (GCC→ACT substitution), without the incorporation of an additional artificial discriminatory mismatch (**Fig. S2c**). *In silico* evaluation demonstrated predicted amplification of a 327 bp product across all *S*. Paratyphi B genomes (100% sensitivity), while excluding the majority of other typhoidal and non-typhoidal serovars (98.35% specificity).

For *S*. Paratyphi C, primers targeting the SPC0571 locus were designed for inclusion in the multiplex panel, generating a predicted 194 bp amplicon. As no discriminatory sequence variants suitable for incorporation at the 3′ primer termini were identified within this locus, primer specificity was limited, resulting in an *in silico* sensitivity of 97.14% and specificity of 99.75%. Predicted cross-amplification was observed with closely related serovars, including *S*. Choleraesuis and *S*. Gallinarum during *in silico* evaluation (**Fig. S2d**). A pan-*Salmonella* primer set targeting the conserved *invA* locus was also designed. Degenerate bases were incorporated to maximize serovar coverage (**Figure S2e**). The resulting 922 bp amplicon demonstrated 100% sensitivity across all *Salmonella* genomes, with no predicted amplification in non-Salmonella genomes.

### *In vitro* Testing, validation, sensitivity and specificity

The performance of the multiplex PCR assay incorporating primers targeting *invA* (pan-*Salmonella*; 922 bp), *staG* (*S*. Typhi; 714 bp), SPA0152 (*S*. Paratyphi A; 539 bp), SPAB_03490 (*S*. Paratyphi B; 327 bp), and SPC0571 (*S*. Paratyphi C; 194 bp) was evaluated using a panel of 30 bacterial isolates (**Table 1**). This panel included five target *Salmonella* serovars Typhi (CT18 and Ty2), Paratyphi A (ATCC 9150), Paratyphi B (clinical strain, KU13854), and. Paratyphi C (RKS4594) along with 17 clinically relevant NTS isolates and eight non-*Salmonella* bacterial species (including *E. coli, K. pneumoniae, P. aeruginosa, A. baumannii, Citrobacter* spp., and *Enterobacter cloacae*).

All target *Salmonella* serovars yielded the expected amplicon profiles corresponding to their respective primer sets, while the *invA* target was consistently amplified across all *Salmonella* isolates, confirming its utility as a pan-*Salmonella* control (**Figure S3a**). The *staG* primer demonstrated specific amplification of *S*. Typhi isolates, whereas SPA0152, SPAB_03490, and SPC0571 primers generated serovar-specific amplicons for *S*. Paratyphi A, *S*. Paratyphi B, and *S*. Paratyphi C, respectively (**Figure S3b–e**). No amplification of serovar-specific targets was observed among non-target *Salmonella* serovars or non-*Salmonella* bacterial pathogens for *staG* and SPA0152, confirming high analytical specificity (**Figure S3b-e**). However, limited cross-reactivity was observed for the SPAB_03490 primer set, with amplification detected in *S*. Weltevreden and *S*. Cerro, and for SPC0571, which showed amplification in *S*. Choleraesuis. These findings are consistent with *in silico* predictions and reflect the absence of fully serovar-exclusive genetic markers for these lineages (**Table 2**).

**Table 2:**
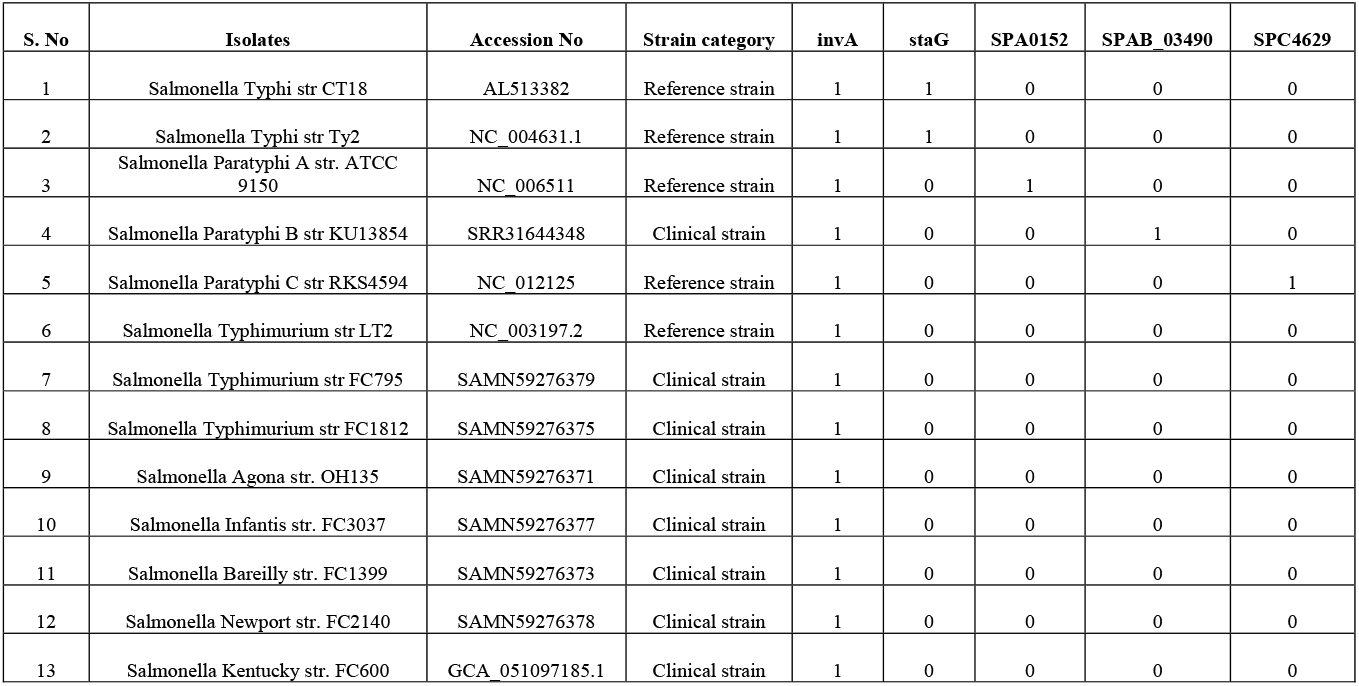
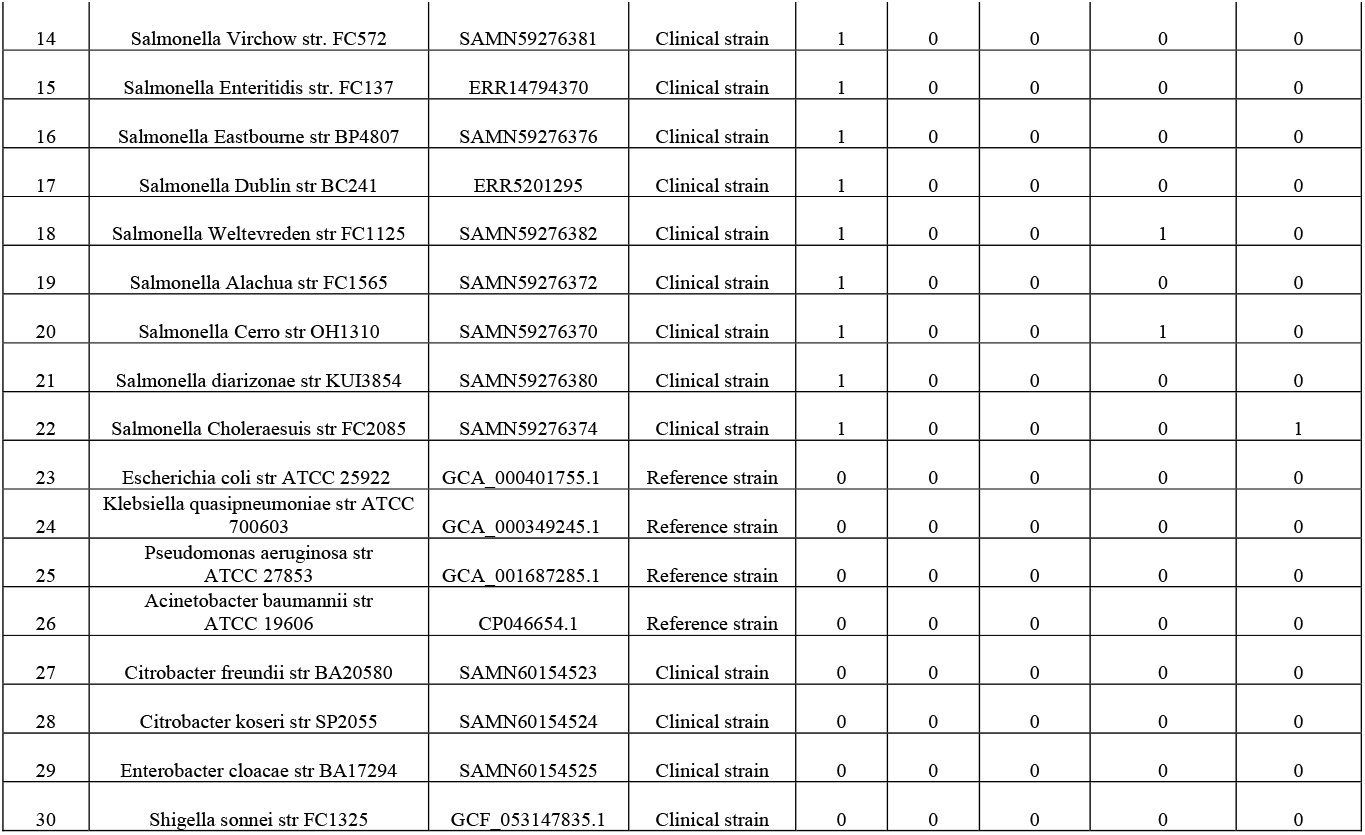
List of Bacterial strains used for in vitro evaluation of the multiplex PCR assay and corresponding amplification profiles.

| S. No | Isolates | Accession No | Strain category | invA | staG | SPA0152 | SPAB_03490 | SPC4629 |
| --- | --- | --- | --- | --- | --- | --- | --- | --- |
| 1 | Salmonella Typhi str CT18 | AL513382 | Reference strain | 1 | 1 | 0 | 0 | 0 |
| 2 | Salmonella Typhi str Ty2 | NC_004631.1 | Reference strain | 1 | 1 | 0 | 0 | 0 |
| 3 | Salmonella Paratyphi A str. ATCC 9150 | NC_006511 | Reference strain | 1 | 0 | 1 | 0 | 0 |
| 4 | Salmonella Paratyphi B str KU13854 | SRR31644348 | Clinical strain | 1 | 0 | 0 | 1 | 0 |
| 5 | Salmonella Paratyphi C str RKS4594 | NC_012125 | Reference strain | 1 | 0 | 0 | 0 | 1 |
| 6 | Salmonella Typhimurium str LT2 | NC_003197.2 | Reference strain | 1 | 0 | 0 | 0 | 0 |
| 7 | Salmonella Typhimurium str FC795 | SAMN59276379 | Clinical strain | 1 | 0 | 0 | 0 | 0 |
| 8 | Salmonella Typhimurium str FC1812 | SAMN59276375 | Clinical strain | 1 | 0 | 0 | 0 | 0 |
| 9 | Salmonella Agona str. OH135 | SAMN59276371 | Clinical strain | 1 | 0 | 0 | 0 | 0 |
| 10 | Salmonella Infantis str. FC3037 | SAMN59276377 | Clinical strain | 1 | 0 | 0 | 0 | 0 |
| 11 | Salmonella Bareilly str. FC1399 | SAMN59276373 | Clinical strain | 1 | 0 | 0 | 0 | 0 |
| 12 | Salmonella Newport str. FC2140 | SAMN59276378 | Clinical strain | 1 | 0 | 0 | 0 | 0 |
| 13 | Salmonella Kentucky str. FC600 | GCA_051097185.1 | Clinical strain | 1 | 0 | 0 | 0 | 0 |
| 14 | Salmonella Virchow str. FC572 | SAMN59276381 | Clinical strain | 1 | 0 | 0 | 0 | 0 |
| 15 | Salmonella Enteritidis str. FC137 | ERR14794370 | Clinical strain | 1 | 0 | 0 | 0 | 0 |
| 16 | Salmonella Eastbourne str BP4807 | SAMN59276376 | Clinical strain | 1 | 0 | 0 | 0 | 0 |
| 17 | Salmonella Dublin str BC241 | ERR5201295 | Clinical strain | 1 | 0 | 0 | 0 | 0 |
| 18 | Salmonella Weltevreden str FC1125 | SAMN59276382 | Clinical strain | 1 | 0 | 0 | 1 | 0 |
| 19 | Salmonella Alachua str FC1565 | SAMN59276372 | Clinical strain | 1 | 0 | 0 | 0 | 0 |
| 20 | Salmonella Cerro str OH1310 | SAMN59276370 | Clinical strain | 1 | 0 | 0 | 1 | 0 |
| 21 | Salmonella diarizonae str KUI3854 | SAMN59276380 | Clinical strain | 1 | 0 | 0 | 0 | 0 |
| 22 | Salmonella Choleraesuis str FC2085 | SAMN59276374 | Clinical strain | 1 | 0 | 0 | 0 | 1 |
| 23 | Escherichia coli str ATCC 25922 | GCA_000401755.1 | Reference strain | 0 | 0 | 0 | 0 | 0 |
| 24 | Klebsiella quasipneumoniae str ATCC 700603 | GCA_000349245.1 | Reference strain | 0 | 0 | 0 | 0 | 0 |
| 25 | Pseudomonas aeruginosa str ATCC 27853 | GCA_001687285.1 | Reference strain | 0 | 0 | 0 | 0 | 0 |
| 26 | Acinetobacter baumannii str ATCC 19606 | CP046654.1 | Reference strain | 0 | 0 | 0 | 0 | 0 |
| 27 | Citrobacter freundii str BA20580 | SAMN60154523 | Clinical strain | 0 | 0 | 0 | 0 | 0 |
| 28 | Citrobacter koseri str SP2055 | SAMN60154524 | Clinical strain | 0 | 0 | 0 | 0 | 0 |
| 29 | Enterobacter cloacae str BA17294 | SAMN60154525 | Clinical strain | 0 | 0 | 0 | 0 | 0 |
| 30 | Shigella sonnei str FC1325 | GCF_053147835.1 | Clinical strain | 0 | 0 | 0 | 0 | 0 |

Despite these minor cross-reactivity events, the multiplex assay accurately identified all target serovars within the tested panel, demonstrating 100% sensitivity for detection of all target serovars within the evaluated isolate panel (**Figure 2**). Importantly, these cross-reactivity events did not affect accurate identification of the target serovars within the multiplex framework. The multiplex design also enabled clear resolution of all amplicons, with distinct band separation across the expected size range.

**Figure 2:**
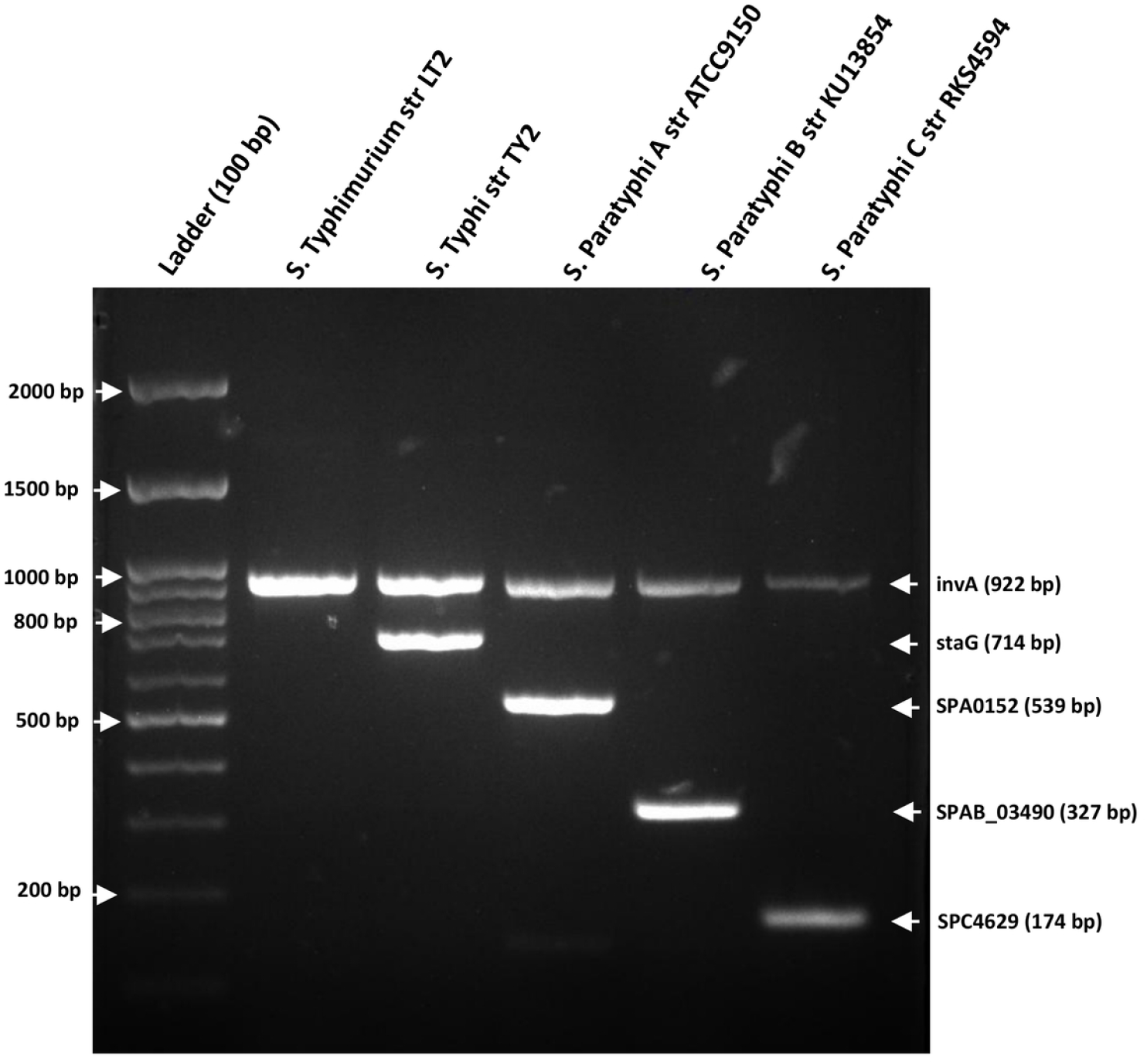
2% agarose gel depicting multiplex PCR amplification profiles of typhoidal and non-typhoidal Salmonella reference strains. Multiplex PCR targeting *invA* (pan-Salmonella; 922 bp), *staG* (*S*. Typhi; 714 bp), SPA0152 (*S*. Paratyphi A; 539 bp), SPAB_03490 (*S*. Paratyphi B; 327 bp), and SPC4629 (*S*. Paratyphi C; 174 bp) was performed using representative reference strains. Lane 1, 100 bp DNA ladder; lane 2, *Salmonella* Typhimurium LT2; lane 3, *S*. Typhi Ty2; lane 4, *S*. Paratyphi A ATCC 9150; lane 5, *S*. Paratyphi B KU13854; lane 6, *S*. Paratyphi C RKS4594. The *invA* amplicon was detected in all Salmonella isolates, whereas serovar-specific amplicons were observed only in their corresponding target strains. Expected amplicon sizes are indicated by arrows on the right margin.

## Discussion

Differentiation of typhoidal *Salmonella* from NTS serovars and other bacterial pathogen is critical for patient management, antimicrobial therapy, and public health surveillance. However, this remains challenging due to the high genomic conservation between typhoidal and non-typhoidal lineages and the extensive diversity within NTS serovars (15). This is particularly relevant in endemic regions, where NTS serovars are increasingly reported as causes of invasive bloodstream infections (16,17). The emergence of *S*. Typhi strains resistant to currently treating antimicrobials further underscores the need for rapid serovar-level identification for appropriate clinical management (31,32). Since blood culture, the reference standard for enteric fever diagnosis, requires 2–7 days for confirmation (18), rapid molecular differentiation of typhoidal serovars has important diagnostic and therapeutic implications (19). In this study, we integrated large-scale comparative genomics with experimental validation to identify robust PCR primers enabling differentiation of all four typhoidal *Salmonella* serovars from NTS serovars.

Our comprehensive *in silico* evaluation of previously published PCR primers revealed significant limitations in the specificity and specificity of many currently used assays. Many classical targets were developed prior to the widespread availability of WGS data and were selected using limited strain collections rather than population-scale genomic evidence (20– 22). Consequently, although several targets demonstrated high predicted sensitivity, their specificity was frequently compromised by cross-reactivity with NTS serovars. This was evident for commonly used *S*. Typhi markers such as *staG* and *tviB*, which were also detected in non-target serovars (6). Similarly, loci commonly used for detection of *S*. Paratyphi A, B, and C were widely distributed across diverse NTS backgrounds (24). To address such limitations, strategies incorporating multiple targets for a single serovar have increasingly been adopted (10,41). More recently, PCR assays targeting typhoidal *Salmonella* have also been described using newly identified targets such as STY4578, STY3279 etc (52). However, these approaches relied primarily on gene-content differences derived from relatively limited genome collections, whereas the present study employed population-scale SNP analysis across 3,239 genomes representing 119 serovars, enabling more comprehensive evaluation of target conservation and cross-reactivity.

Guided by these observations, we initially employed a pangenome-based strategy to identify serovar-specific targets. Consistent with previous studies, our analysis confirmed that *Salmonella enterica* possesses a largely closed pangenome with extensive sharing of core genes across serovars (23,24) Consequently, truly unique genes exclusively conserved within individual typhoidal serovars were rare (53). Moreover, several putative serovar-specific genes identified during pangenome analysis were found to represent misannotated pseudogenes rather than true lineage-specific targets. This is particularly relevant for host-adapted typhoidal serovars, which have undergone reductive genome evolution associated with pseudogene accumulation (25). These findings highlight an important limitation of automated pangenome pipelines and emphasize the need for sequence-level validation of unique genes prior to diagnostic assay development based on pangenome analysis (54). Our findings demonstrate that reliance on gene presence alone is insufficient for robust serovar discrimination and instead support SNP-based primer design within conserved loci. Using MAMA-PCR principles, discriminatory SNPs were exploited by incorporating deliberate 3′ mismatches to enhance allele-specific amplification (26). This enabled reliable differentiation of closely related typhoidal serovars using lineage-specific SNPs, an approach previously shown to improve discrimination among highly conserved *Salmonella* lineages (27,28). The resulting primer sets achieved predicted specificities of 99–100%, demonstrating that SNP-based discrimination within conserved genes provides a robust framework for molecular differentiation of typhoidal *Salmonella*.

Importantly, the feasibility of identifying fully exclusive targets varied across serovars (23,29). While *S*. Typhi and *S*. Paratyphi C permitted highly discriminatory SNP-based designs, *S*. Paratyphi A and *S*. Paratyphi B required optimization strategies balancing sensitivity and specificity (30). *In vitro* testing showed concordance with *in silico* predictions and improved specificity compared with previously described assays, particularly for *S*. Paratyphi A and *S*. Paratyphi B. Limited cross-reactivity observed for *S*. Paratyphi B and *S*. Paratyphi C targets reflects the absence of fully exclusive markers within the genus. However, the implicated serovars (*S*. Weltevreden, *S*. Cerro, and *S*. Choleraesuis) are infrequently associated with invasive bloodstream infections in typhoid-endemic settings, reducing the practical clinical impact of these findings (7). In addition, incorporation of the pan-*Salmonella invA* target provides an important interpretive advantage, as amplification of *invA* in the absence of serovar-specific products indicates probable NTS infection rather than assay failure (49). Overall, the multiplex assay enabled simultaneous differentiation of all four typhoidal serovars and demonstrated strong potential for diagnostic and surveillance applications in endemic settings.

This study has several limitations. Although *in silico* evaluation was performed using a large and globally representative genome dataset, *in vitro* validation included a limited number of reference and clinical isolates. Rare NTS lineages predicted to share sequence similarity with selected targets were not represented experimentally. In addition, assay performance was evaluated using cultured isolates, and further studies using primary clinical specimens and geographically diverse collections will be necessary to confirm field performance. Future work should also evaluate applicability to blood culture workflows and alternative formats such as real-time or isothermal amplification assays.

## Conclusion

In conclusion, this study demonstrates that genomics-informed, SNP-based primer design enables reliable differentiation of all four typhoidal *Salmonella* serovars despite extensive genomic conservation with non-typhoidal lineages. Integration of population-scale *in silico* analysis with experimental validation improved diagnostic specificity without compromising analytical sensitivity, overcoming key limitations of conventional gene-based assays. The strong concordance between *in silico* and *in vitro* performance supports the utility of this multiplex assay for diagnostic and surveillance applications in endemic settings. More broadly, this approach provides a scalable framework for molecular assay development in other genetically conserved bacterial pathogens.

## Acknowledgment

We express our gratitude to the Department of Clinical Microbiology, Christian Medical College, Vellore, for providing the essential facilities and support that made this study possible. We are also grateful to Ms. Praveena Jeslin, Ms. Baby Abirami Shankar, Ms. Tharani Priya T and Ms. Devi Shree for their valuable assistance in stock culture maintenance.

## Ethical Clearance

Institutional Review Board (IRB) of Christian Medical College (CMC), Vellore, India gave ethical approval for this work vide IRB Min no. 16106 dated 28.02.2024. As the required data had been collected as part of the standard of care for diagnosis, informed consent waiver was granted by the IRB committee Christian Medical College, Vellore.

## Contributors

Conceptualization: JJJ and BV

Methodology and Investigation: JJJ, PT, PS, KBS, SR, AV, and MPT

Data analysis: JJJ and PT

Visualization: JJJ and PT

Funding acquisition: BV

Project administration: JJJ, NP, AN

Supervision: JJ, BV and KW

Original draft: JJJ, PT

Review & editing: JJ, BV and KW

## Patient consent for publication

Not required.

## Data availability statement

Raw read data were deposited in the European Nucleotide Archive (ENA) under project accession number: PRJNA1463158. The individual sample accession numbers are listed in Table S1 & Table 2

## Figure legends

**Figure S1: Pangenome structure and distribution of lineage-specific core genes across Salmonella serovar groups. (a)** Pie chart showing the distribution of the Salmonella pangenome into core genes (present in ≥99% of genomes), soft-core genes (95–99%), shell genes (15–95%), and cloud genes (<15%) identified using Panaroo **(b)** Dot plot generated using Twilight pangenome clustering showing the distribution of lineage-specific core genes across non-typhoidal Salmonella (NTS), S. Paratyphi B (SPB), S. Paratyphi A (SPA), S. Typhi (Typhi), S. Paratyphi C (SPC), and genomes with unresolved serovar assignment (Unknown). Each dot represents an individual genome, and horizontal black lines indicate median values for each group. *S*. Typhi showed the highest number of lineage-specific core genes, followed by S. Paratyphi A and *S*. Paratyphi C, whereas NTS and *S*. Paratyphi B genomes showed few or no lineage-specific core genes.

**Figure S2: Multiple sequence alignment of primer binding regions across representative Salmonella genomes.** Primer binding regions corresponding to the (a) *staG* (S. Typhi), (b) SPA0152 (*S*. Paratyphi A), (c) SPAB_03490 (*S*. Paratyphi B), (d) SPC4629 (*S*. Paratyphi C) and (e) *invA* (pan-Salmonella), targets were aligned against representative Salmonella genomes using Bioedit v7.1. Primer sequences are shown in bold above the aligned genomic regions with corresponding GenBank accession numbers. Conserved nucleotides are represented by identical colour-coded blocks (A, green; T, red; G, yellow; C, blue), whereas mismatches relative to the primer sequence are shown as variable positions.

**Figure S3: 2% agarose gel electrophoresis showing analytical specificity of individual PCR primer sets against Salmonella and non-Salmonella bacterial isolates.** Singleplex PCR assays were performed using primers targeting (A) *invA* (922 bp), (B) staG (714 bp), (C) SPA0152 (539 bp), (D) SPAB_03490 (327 bp), and (E) SPC4629 (174 bp). M, 100 bp DNA ladder. Lane assignments: 1, *Salmonella* Typhi str. CT18; 2, *Salmonella* Typhi str. Ty2; 3, *Salmonella* Paratyphi A str. ATCC 9150; 4, *Salmonella* Paratyphi B str. KU13854; 5, *Salmonella* Paratyphi C str. RKS4594; 6, *Salmonella* Typhimurium str. LT2; 7, *Salmonella* Typhimurium str. FC795; 8, *Salmonella* Typhimurium str. FC1812; 9, *Salmonella* Agona str. OH135; 10, *Salmonella* Infantis str. FC3037; 11, *Salmonella* Bareilly str. FC1399; 12, *Salmonella* Newport str. FC2140; 13, *Salmonella* Kentucky str. FC600; 14, *Salmonella* Virchow str. FC572; 15, *Salmonella* Enteritidis str. FC572; 16, *Salmonella* Eastbourne str. BP4807; 17, *Salmonella* Dublin str. BC241; 18, *Salmonella* Weltevreden str. FC1125; 19, *Salmonella* Alachua str. FC1565; 20, *Salmonella* Cerro str. OH1310; 21, *Salmonella* diarizonae str. KU13854; 22, *Salmonella* Choleraesuis str. FC2085; 23, *Escherichia coli* str. ATCC 25922; 24, *Klebsiella quasipneumoniae* str. ATCC 700603; 25, *Pseudomonas aeruginosa* str. ATCC 27853; 26, *Acinetobacter baumannii* str. ATCC 19606; 27, *Citrobacter freundii* str. BA20580; 28, *Citrobacter koseri* str. SP2055; 29, *Enterobacter cloacae* str. BA17294; 30, *Shigella sonnei* str. FC1325; 31, no-template negative control. The *invA* primer set amplified all Salmonella isolates without amplification in non-Salmonella organisms, confirming genus-level specificity. The *staG*, SPA0152, SPAB_03490, and SPC4629 primer sets produced amplicons specific for S. Typhi, S. Paratyphi A, S. Paratyphi B, and S. Paratyphi C, respectively. Limited cross-reactivity was observed for SPAB_03490 with S. Weltevreden and S. Cerro, and for SPC4629 with S. Choleraesuis, consistent with in silico predictions.

